# Variability of dynamic patterns of cortical excitability in schizophrenia: A test-retest TMS-EEG study

**DOI:** 10.1101/2020.06.24.170019

**Authors:** Jodie Naim-Feil, Abraham Peled, Dominik Freche, Shmuel Hess, Alexander Grinshpoon, Nava Levit-Binnun, Elisha Moses

## Abstract

**Background:** Altered stimuli processing is a key feature of schizophrenia. Application of concurrent transcranial magnetic stimulation (TMS) with electroencephalography (TMS-EEG) is an effective stimulus which allows direct measurement of the cortical response within a millisecond time resolution. Test-retest TMS-EEG studies present evidence of high reproducibility in healthy controls (HC), however, this stability of response has not been examined in schizophrenia.

**Objective:** The current study maps TMS-evoked patterns of cortical excitability in schizophrenia and examine whether these cortical patterns are amenable to change as symptoms of schizophrenia improve.

**Methods:** One hundred single-pulse TMS and 100 sham pulses were applied to frontal regions of 19 schizophrenia in-patients and 26 HC while electroencephalography data were simultaneously acquired. Medication and schizophrenia symptoms were reported. This protocol was repeated across three sessions (1 week apart) for each participant. The TMS-evoked cortical response of each participant was averaged and compared between groups.

**Results:** Schizophrenia patients showed reduced cortical excitability at early time-windows and increased excitability at later time-windows. Increased excitability at later windows was associated with heightened symptom severity. Schizophrenia patients showed an increased variability in cortical response over sessions relative to HC. Increased change in cortical response from session 1 to session 3 correlated with symptom improvement.

**Conclusions:** Schizophrenia patients presented with abnormal patterns of cortical excitability when processing TMS stimuli. These dynamic patterns of cortical response were amenable to change as symptoms of schizophrenia improved. Further research into electrophysiological biomarkers of symptom improvement is hoped to improve current diagnosis and treatment models of schizophrenia.

## Introduction

Schizophrenia (SCH) is a chronic and debilitating disorder which is characterised by a number of heterogenous symptoms. Both positive and negative symptoms of SCH were traditionally viewed as stable and thought to increase over time as the disease progresses [1]. Recent experimental studies have demonstrated symptom improvement in response to various intervention approaches [2], which are however prone to relapse [3]. Identification of clinically useful predictors of symptom improvement may offer new insight into treatment responsiveness and relapse prevention. In particular, disturbance of stimuli processing (such as impaired sensory gating) is considered a central feature of SCH and is commonly associated with exacerbated symptoms [4–6]. Sensory gating is associated with the ability to filter out irrelevant sensory stimuli and protect against flooding higher order cortical areas with information [7,8]. The current study is designed to explore whether disrupted stimuli processing can improve over time in SCH patients receiving treatment and whether these changes in stimuli processing are associated with symptom improvement.

The combination of transcranial magnetic stimulation [9,10] with electroencephalography (TMS-EEG) is an effective stimulus with a direct physiological measure of the local and interconnected brain response with millisecond time resolution [11,12]. A significant advantage of TMS-EEG within a population with SCH is that it is direct, unmitigated [13,14], and does not require active participation of the participants [15–19]. TMS-EEG is therefore emerging as a promising technique for mapping stimuli processing in SCH [13].

To date, use of the single-pulse TMS paradigm has mainly focused on the early cortical response to a TMS perturbation (first 300ms following TMS) when applied to various brain regions, such as the motor, premotor, prefrontal and parietal regions [15,20–22]. Compelling evidence of widespread *reduced* excitability, altered patterns of propagation, and dampened oscillatory response to TMS was reported in SCH patients. However, Frantseva et al (2014) examined late components of TMS-evoked cortical excitability in SCH patients, and demonstrated *increased* excitability at the later windows (200ms to 900ms). The increased activity at later time windows contrasts with the reduced activity at early time windows, and highlights the importance of exploring the dynamic brain processes in adequately quantifying the temporal patterns of stimuli response in SCH patients. Interestingly, both early and late components of TEP have separately been previously linked to symptom severity [20,23]. The current study aims to characterise how dynamic patterns of TMS-evoked cortical response evolve over an extended time course in SCH patients.

Once the dynamic patterns of cortical response have been quantified, the present study will explore the reproducibility of these patterns and ascertain whether changes in cortical response over time relate to treatment response. Examining the parallel changes over a significant period in SCH patients of both symptoms and response to TMS necessitates a test-retest design. Test-retest of TMS-EEG typically involves precise repetition of a particular design paradigm with a waiting interval on the order of a week. This approach has only been utilized by a small number of TMS-EEG studies, mainly to demonstrate the reliability of concurrent TMS-EEG in healthy controls [24–26]. High reproducibility between sessions has been observed in healthy controls when coil location and coil orientation are kept consistent and is robust to varying TMS intensities once a relatively low intensity threshold is exceeded [27].

In this study we apply the test-retest approach within a treatment-receiving group of SCH patients, examining whether SCH patients are more variable between sessions in the response to single pulse TMS stimuli compared to healthy controls (HC). We anticipate that changes in cortical response will correlate with symptom improvement, which will allow us to identify those time windows that are most amenable to change (over sessions) as symptoms improve. If this occurs, we will have identified a novel method for the detection of electrophysiological biomarkers of symptom improvement in SCH. To our knowledge this is the first study to investigate the relationship between the reproducibility of TMS-evoked cortical response in SCH patients and symptom improvement.

We thus define three major objectives: (1) Apply single-pulse frontal TMS-EEG to characterize the dynamic patterns of cortical response to TMS stimuli in SCH patients compared to HC over an extended time-course. (2) Use a test-retest approach to ascertain whether patterns of TMS-evoked cortical response are more variable over sessions across SCH patients relative to HC. (3) Examine whether, as symptoms of SCH improve over time, a change in the TMS-evoked cortical response over sessions will be detected and whether this increased variability in response will be associated with the symptom improvement.

## Methods and Materials

The study was approved by the Shaar Menashe Mental Health Center Institutional Review Board. Participants were reimbursed the equivalence of 35 US dollars for participating in each of the TMS-EEG sessions.

### Subjects

Thirty in-unit patients at Shaar Menashe Mental Health Center who met the criteria for DSM IV-TR Schizophrenia [28] were recruited. SCH patients presenting with a history of neurology disorders, psychiatric comorbidity and drug abuse were excluded from the study. Five SCH datasets were excluded due to lack of completion of all three sessions of active TMS, a further four datasets were excluded due to lack of completion of all three sessions of sham TMS and another two datasets excluded due to excessive muscle movement.

Therefore, for the present study, data from nineteen SCH patients were analysed. This sample size is consistent with the prevalence of high dropout rates by SCH patients in experimental trials which require multiple sessions [29]. A trained psychiatrist conducted the baseline screening of SCH patients and administered the Scale for the Assessment of Positive Symptoms (SAPS) and the Scale for the Assessment of Negative Symptoms (SANS) to quantify clinical symptoms of SCH (30, 31).

Thirty healthy controls (HC) without any previous or current history of psychiatric illness, drug abuse or head injury were recruited through local advertisements. Two HC datasets were excluded due to excessive muscle movement and a further two datasets were excluded due to lack of completion of all three sessions of sham TMS. Twenty-six HC datasets were analysed.

All participants completed a general demographic questionnaire and TMS safety questionnaire [32]. Demographic and clinical data are described in Table 1.

**Table 1.**
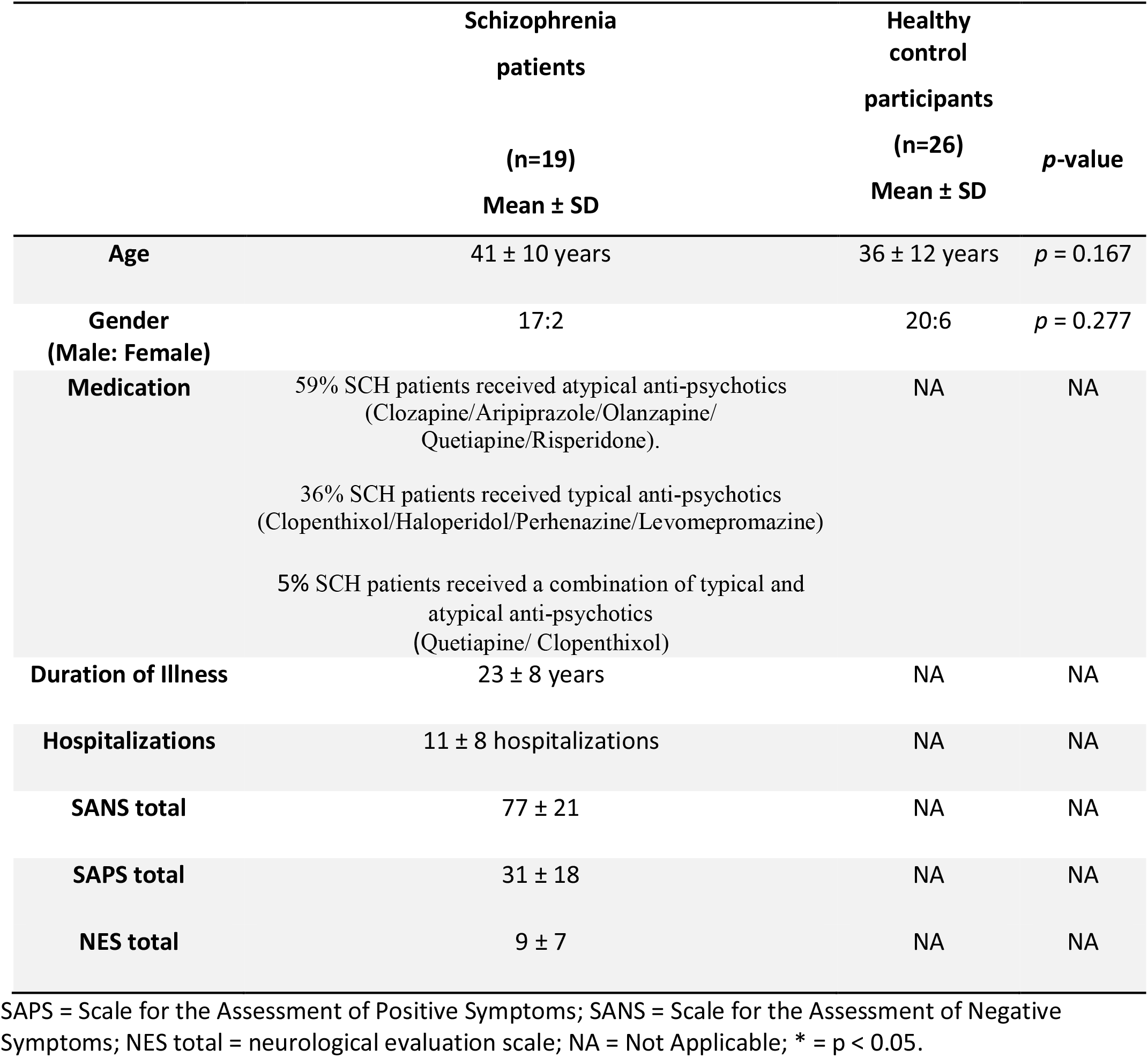
Demographic and clinical data for schizophrenia patients and healthy control participants.

|  | Schizophrenia<br>patients<br>(n=19)<br>Mean ± SD | Healthy<br>control<br>participants<br>(n=26)<br>Mean ± SD | p-value |
| --- | --- | --- | --- |
| **Age** | 41 ± 10 years | 36 ± 12 years | p = 0.167 |
| **Gender**<br>**(Male: Female)** | 17:2 | 20:6 | p = 0.277 |
| **Medication** | 59% SCH patients received atypical anti-psychotics<br>(Clozapine/Aripiprazole/Olanzapine/<br>Quetiapine/Risperidone).<br>36% SCH patients received typical anti-psychotics<br>(Clopenthixol/Haloperidol/Perhenazine/Levomepromazine)<br>5% SCH patients received a combination of typical and<br>atypical anti-psychotics<br>(Quetiapine/ Clopenthixol) | NA | NA |
| **Duration of Illness** | 23 ± 8 years | NA | NA |
| **Hospitalizations** | 11 ± 8 hospitalizations | NA | NA |
| **SANS total** | 77 ± 21 | NA | NA |
| **SAPS total** | 31 ± 18 | NA | NA |
| **NES total** | 9 ± 7 | NA | NA |

SAPS = Scale for the Assessment of Positive Symptoms; SANS = Scale for the Assessment of Negative Symptoms; NES total = neurological evaluation scale; NA = Not Applicable; \* = p < 0.05.

### Experimental design

A BioSemi ActiveTwo EEG measurement system (BioSemi, Amsterdam, The Netherlands) was used with a head-cap holding 64 Ag-AgCl active electrodes and digitized at 1024Hz. To maintain cap placement consistency between sessions, the same EEG cap was used for all sessions with the exact cap location measured and placed accordingly. The psychiatrist then examined three minutes of baseline EEG recordings for evidence of epileptic activity (an exclusion criteria).

Each measurement comprised of 100 single TMS pulses delivered to the frontal cortex of participants (electrode FC2) at 3 seconds intervals by a Magstim Rapid 2 (Magstim Co., Dyfed, UK) at 80% intensity via a figure-8 coil. EEG was used to measure the cortical response. All active TMS sessions were preceded by a control SHAM session (same protocol) with the figure-8 coil placed on its side (edge touching the electrode). A custom-built TMS-stand was used to reduce variability between sessions. The exact coil position/orientation with respect to the FC2 electrode was measured and this coil placement was replicated in each session. To reduce auditory interference, participants listened to white noise through headphones during the sessions.

This exact procedure was conducted three times, at the same time of day, with one week between sessions (Figure 1.). Participants reported their sleep/coffee/cigarette intake and were asked to keep their intake consistent on testing days.

**Figure 1.**
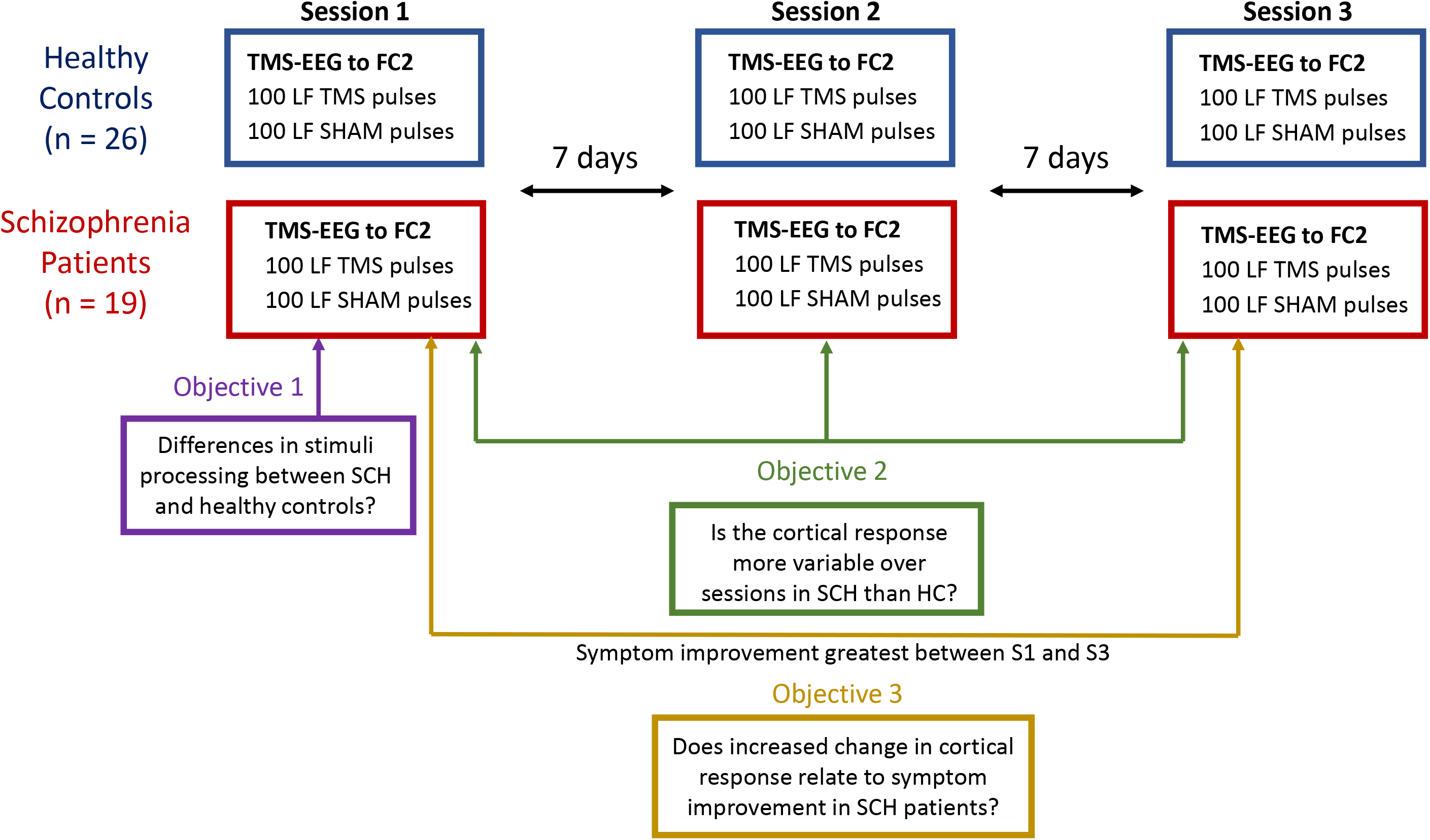
Study procedure timeline. Both schizophrenia patients (SCH: n = 19) and healthy controls (HC: n = 26) participated in all three sessions of concurrent transcranial magnetic stimulation and electroencephalography (TMS-EEG). Prior to testing, participants completed a general demographic and TMS safety questionnaire. This was followed by three testing sessions (a week between each session) in which 100 low frequency pulses of frontal TMS was applied to the FC2 electrode and EEG measure the cortical response. All participants were administered active TMS which were preceded by a control SHAM session.

### Electroencephalography (EEG) processing

EEG recordings were processed offline using the EEGLAB open source toolbox [33] and inhouse Matlab scripts (The MathWorks, Inc, Natick, MA). The Supplementary Materials describes all EEG pre-processing steps as well as provides graphs of TMS-evoked potentials for each electrode (see Figure S1A. and Figure S1B.). The TMS artifact typically dominates the initial response signal for about 100 milliseconds. We applied a newly developed artifact removal algorithm that can easily be automated and produces markedly clean EEG signals over a period beginning as early as 5 milliseconds following the TMS pulse [34].

Based on the dynamic approach described in our previous publication, each epoch was averaged over trials and then divided into six time-windows with respect to the TMS pulse, with Baseline at −850ms to −550ms and Windows 1 through 6 at 50 to 350ms, 350-650ms, 650-850ms, 850-1200ms, 1200-1500ms and 1500-1800ms respectively [35].

### TMS-evoked Potential (TEP) Analysis

For TEP comparisons the squared mean voltage (referred to as the mean amplitude) for each time-window was calculated. To evaluate the *change over time* in TEP between sessions, we calculated the mean amplitude at Session 1 minus the mean amplitude at Session 3 for each participant, and normalised by the mean of Session 1 and Session 3. These specific sessions (i.e. the difference between Session 1 and Session 3) were chosen as symptom improvement (improved scoring on the SAPS/SANS) was greatest between these sessions.

## Statistical analysis

For all statistical tests, an alpha value of *p* < 0.05 was applied. Data analyses were performed using SPSS for Windows, version 22 and in-house Matlab scripts. The basic demographics of SCH and HC were compared using chi-squared tests for categorical and t-tests for continuous variables (Table 1). In cases where Levene’s test of equal variances were violated, the Welch *p*-value was reported. Following the electrode-wise statistical tests, multiple comparisons were accounted for by use an FDR-controlling procedure with a significance threshold of α = 5%.

The cortical responses were examined over an extended time-window (50-1800 ms). Based on our previous work, the 64 electrodes were divided into fourteen representative brain regions (see Supplementary Materials for details, specifically see Figure S2). From these 14 representative brain regions, 7 regions of interest (across the Prefrontal Cortex, Sensory-Motor and Parietal Regions) to reduce the number comparisons were chosen based on 1.

Topological maps of change over time (Figure 2.) and 2. According to their neurobiological role in stimulus processing. Participant average TEPs across each of the time windows, averaged over each of the brain regions, were extracted. Analysis of variance (ANOVA) was applied to examine whether there are significant differences between the groups in response to TMS at Session 1. Pearson’s correlation was applied to examine the relationship between the TEP response and symptoms of SCH. Variability over sessions of TMS-evoked cortical response in SCH patients relative to HC was evaluated using the Standard Error of Measurement (SEM: see AERA et al., 1985). This is a useful approach for observing variability over sessions [25,36], whereas the larger the SEM, the more variable the score across the sessions (as detailed in Supplementary Materials). The SEM was applied to the mean TEP amplitude of the three sessions for each of the 64 electrodes for each group (SCH and HC). SEM values were normalised by the mean TEP amplitude of the three sessions (for each group) and extracted for each of the time-windows.

**Figure 2.**
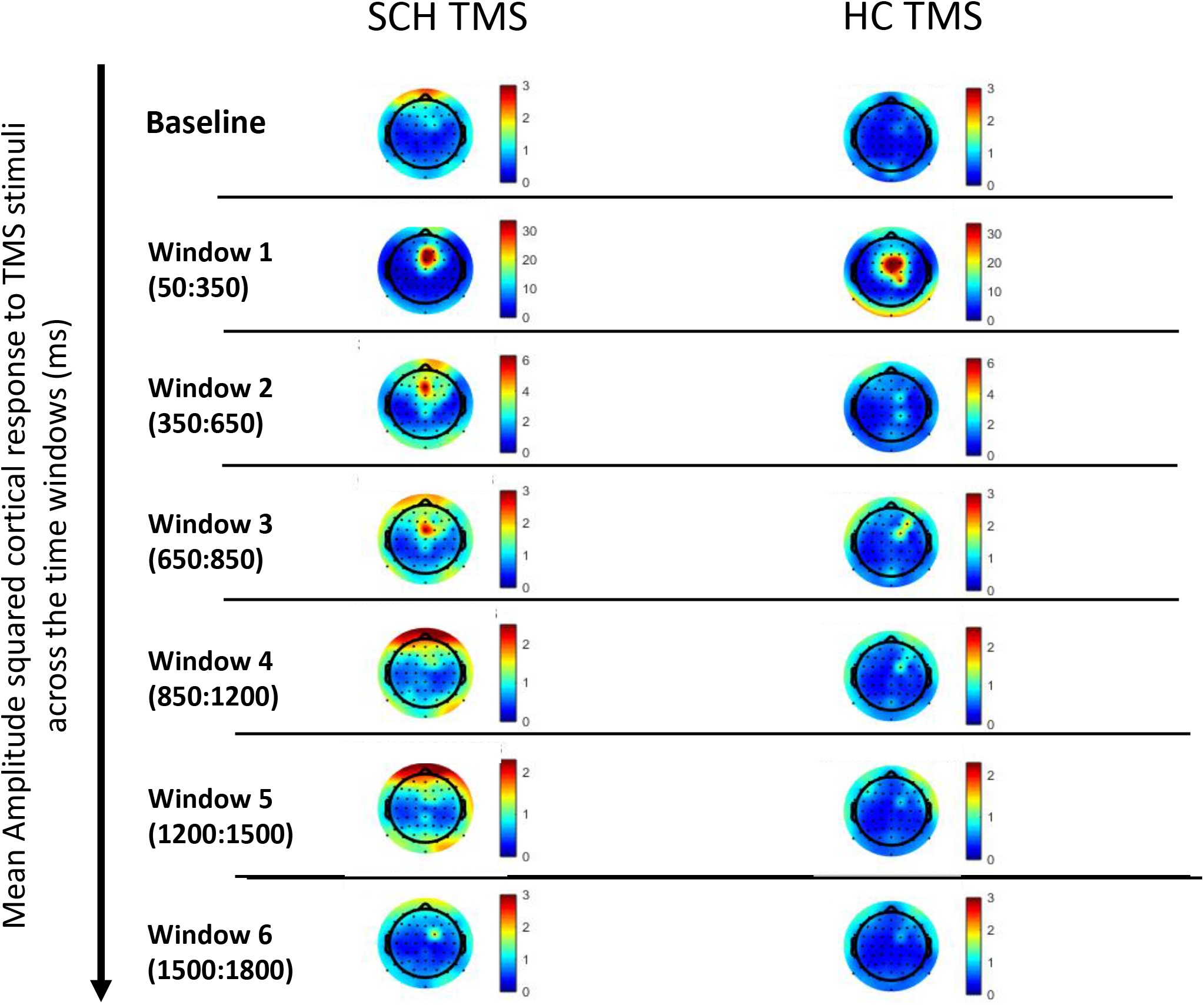
Topographical representation of the averaged dynamic patterns of cortical response to TMS stimuli applied to the frontal cortex in schizophrenia patients (SCH: n = 19) and Healthy Controls (HC: n = 26) across the 64 electrodes. The cortical response is measured at baseline (−850:-550ms pre-pulse), Window 1 (50:350ms), Window 2 (350:650ms), Window 3 (650:850ms), Window 4 (850:1200ms), Window 5 (1200:1500ms) and Window 6 (1500:1800ms). The colorbar varies from blue (minimal activation) to red (maximal activation) and reflects the maximum mean amplitude squared values averaged across groups for each timepoint. Note the change of colorbar units between time windows.

Changes between week 1 to week 3 was then statistically quantified by examining the absolute values of the change in TEP over sessions (from Session 1 to Session 3) averaged across each brain region, and across each time window for each of the participants.

Most SCH patients showed either improvement or no change in negative symptoms (NEG: n = 18) and likewise for positive symptoms (POS: n = 13). There was significant overlap between the groups, with all participants within the POS (n=13) also meeting the criteria for the NEG (n=18) subgroup. This subset was used to explore whether reproducibility of cortical response relates to symptom improvement. Pearson’s correlation was applied to examine whether the TEP change over sessions correlated with improvement of SCH symptom (from Session 1 to Session 3).

## Results

### TMS-evoked cortical response at Session 1

Figure 2. maps the dynamic patterns of cortical response to the TMS stimuli in the 64 electrodes, averaged across SCH patients and across HC. Window 1 demonstrates, in contrast, an attenuation in response of SCH compared to HC. While, in contrast, increased cortical response is observed in SCH in the later time windows, from Window 2 onwards. An appearance of increased frontal activity at baseline in SCH (compared to HC) is also present.

The cortical response averaged over 7 representative brain regions is detailed in the Supplementary Materials (see Figure S4.).

Figure 3A shows, for Session 1, the location of significant differences in response between SCH patients and HC, after grouping of the 64 electrodes and extracting the seven regions of interest. At Window 1 SCH patients presented with reduced cortical activity relative to HC in the left sensory-motor region. At the later windows (Windows 4 – 5) SCH patients showed increased cortical activity across the frontal and parietal regions in response to the TMS stimuli compared to HC (see Table 2).

**Figure 3.**
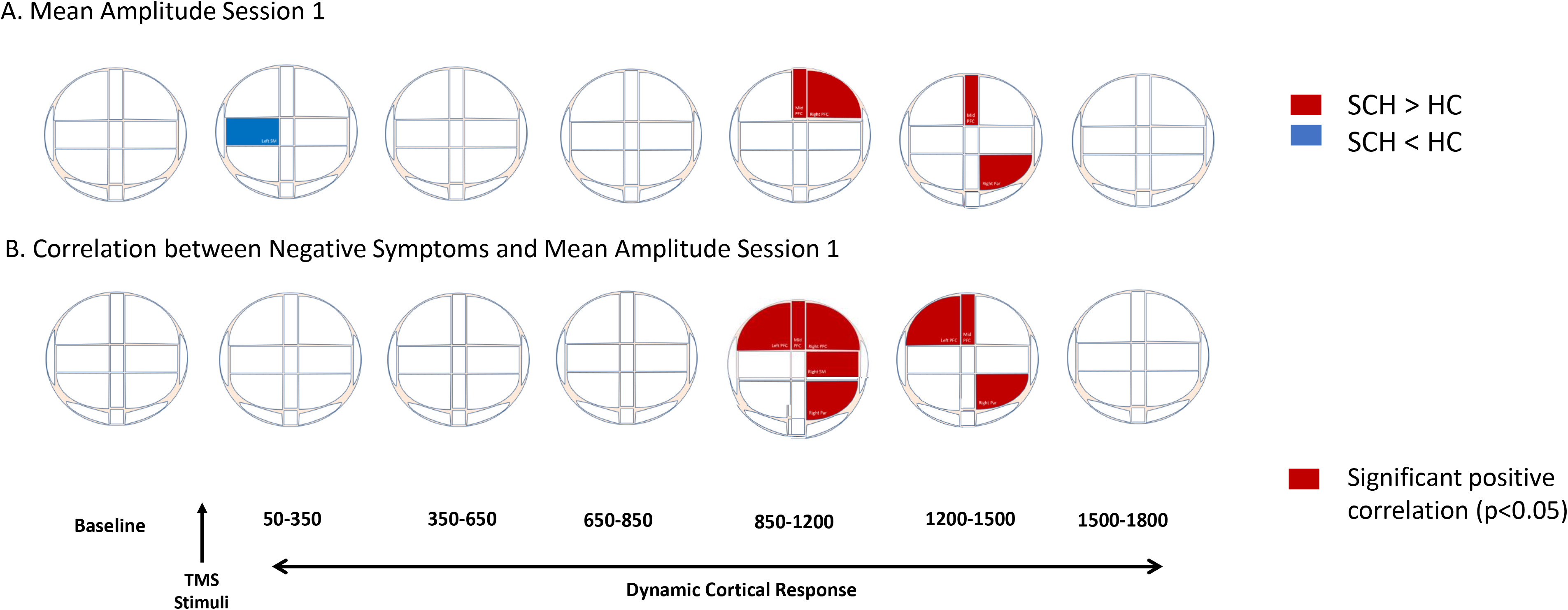
Significant differences in dynamic patterns of cortical response to frontal TMS stimuli at session 1. **Figure 3A.** Shows the regions of statistically significant differences in cortical conductivity for schizophrenia (SCH) patients and healthy controls (HC). Red regions reflect areas in which SCH is significantly higher than HC, and blue regions represent areas in which SCH is significantly lower than HC. **Figure 3B.** Demonstrates the locations of significant correlation between the mean amplitude squared at Session 1 and negative symptoms of SCH at Session 1. There were no significant correlations observed between mean amplitude squared at Session 1 and positive symptoms of SCH at Session 1. All significance values are set at an alpha value of *(p* < 0.05) and only positive correlations were found between mean amplitude and symptoms of SCH indicating that there is a relationship between the increased cortical excitability and severity of clinical symptoms of SCH.

**Table 2.** Significant differences in cortical response to the TMS stimuli, presented as mean amplitude squared (and standard deviation) for each of the brain regions across each of the time windows for schizophrenia patients and healthy controls at Session 1. To account for the multiple comparison, we have presented the original *p*-value as well as the FDR-adjusted (for α = 5%) *p*-value. Only data which differed significantly between the groups (following the FDR-adjustment) is presented in the table.

| | Healthy controls<br>(n=26, $\mu V$)<br>Mean | Healthy controls<br>(n=26, $\mu V$)<br>STD | SCH patients<br>(n=19, $\mu V$)<br>Mean | SCH patients<br>(n=19, $\mu V$)<br>STD | F-value<br>$df = (1,49)$ | Original<br>$p$-value | FDR<br>adjusted<br>$p$-value |
| --- | --- | --- | --- | --- | --- | --- | --- |
| **Window 1** |  |  |  |  |  |  |  |
| SM Left | 9.57 | 8.37 | 4.18 | 3.06 | 7.968 | 0.009 | 0.048 |
| **Window 4** |  |  |  |  |  |  |  |
| PFC Right | 0.7615 | 0.69491 | 1.3318 | 0.94698 | 5.441 | 0.024 | 0.042 |
| PFC Mid | 0.55 | 0.38 | 1.47 | 1.29 | 3.528 | 0.004 | 0.028 |
| **Window 5** |  |  |  |  |  |  |  |
| PFC Mid | 0.43 | 0.29 | 1.41 | 1.53 | 8.210 | 0.01 | 0.035 |
| PAR Right | 0.45 | 0.29 | 0.76 | 0.56 | 6.547 | 0.017 | 0.040 |

Figure 3B demonstrates, for Session 1, the location of significant correlations between the cortical response to TMS and negative symptom severity. Significant positive correlations were observed between increased cortical activity at the later windows (across Windows 4 and 5) and severity of negative symptoms across the frontal, sensory-motor and parietal regions in the SCH patients. There were no significant correlations between the cortical response to TMS and positive symptom severity. No significant correlations were observed at baseline or across the early window (Window 1) or late window (Window 6) for symptoms (significant correlation data detailed in Table 3.).

### Changes in TMS-evoked cortical response between Sessions

Next, the SEM was calculated to ascertain whether SCH patients present overall with increased variability in cortical response over all sessions compared to HC. Figure 4 provides a topographical representation of the SEM values across electrodes and demonstrates increased SEM (indicating greater variability) across all time windows in SCH patients relative to HC (see Figure S3 of Supplementary Materials for SEM values across each electrode).

**Figure 4.**
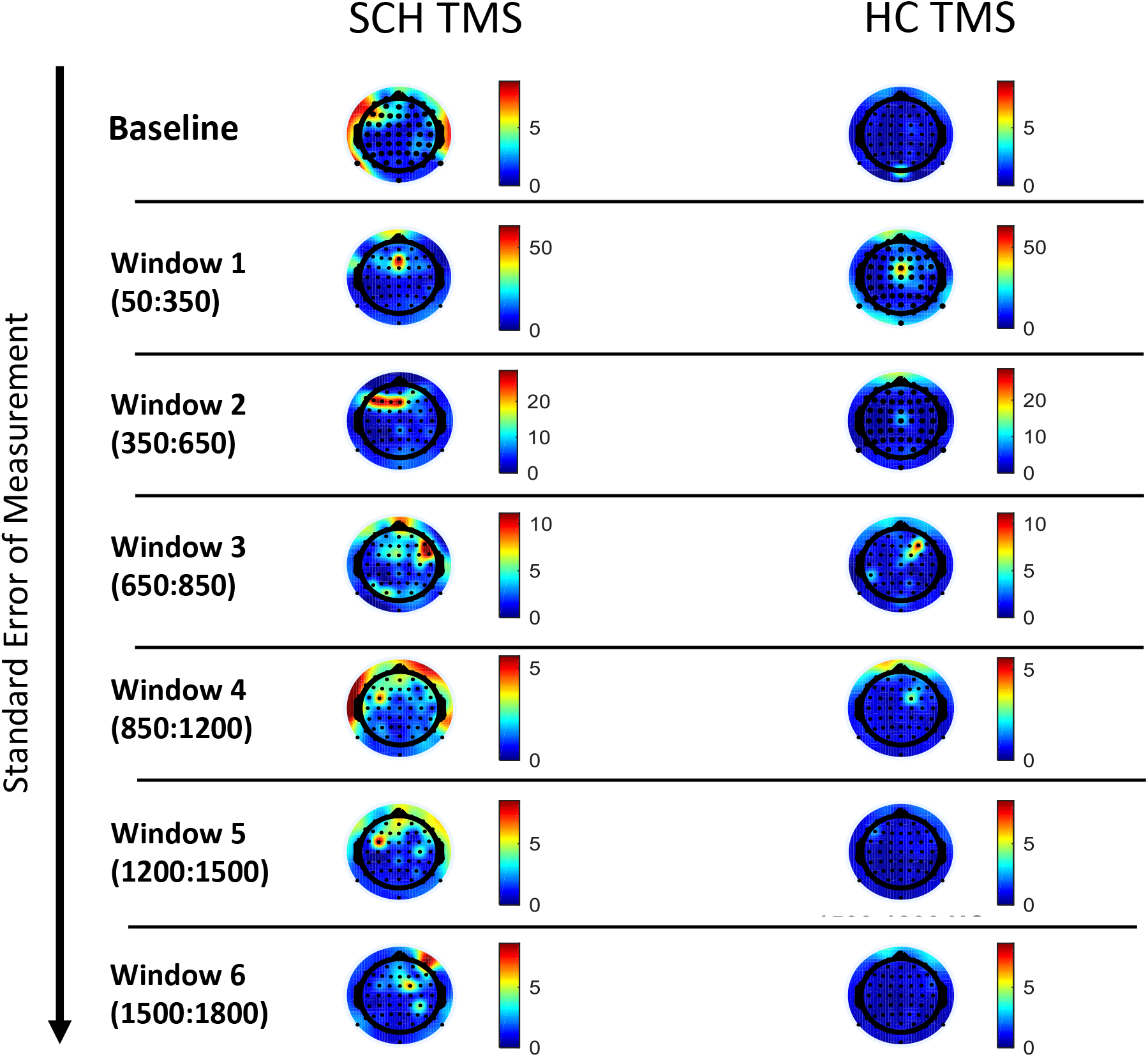
Topographical representations of the Standard Error of Measurement for schizophrenia (SCH) patients and healthy controls (HC). The topographical map shows the standard error of measurement (SEM) values calculated across the mean TEP amplitude of the three sessions for the schizophrenia patients (SCH) compared to healthy controls (HC) for each electrode. The SEM of the three sessions are normalised by the mean amplitude of the three sessions. The larger the SEM the *less reliability* between the three sessions. The topographical map demonstrates the location of the SEM amplitudes across the electrodes and illustrates how the SEM values change over the different time windows. SCH patients show predominantly increased SEM values compared to HC across the baseline, Window 2, 3, 4, 5 & 6. Therefore, SCH appears to show a more variable response to the TMS pulse across the 3 sessions than HC.

**Table 3.**
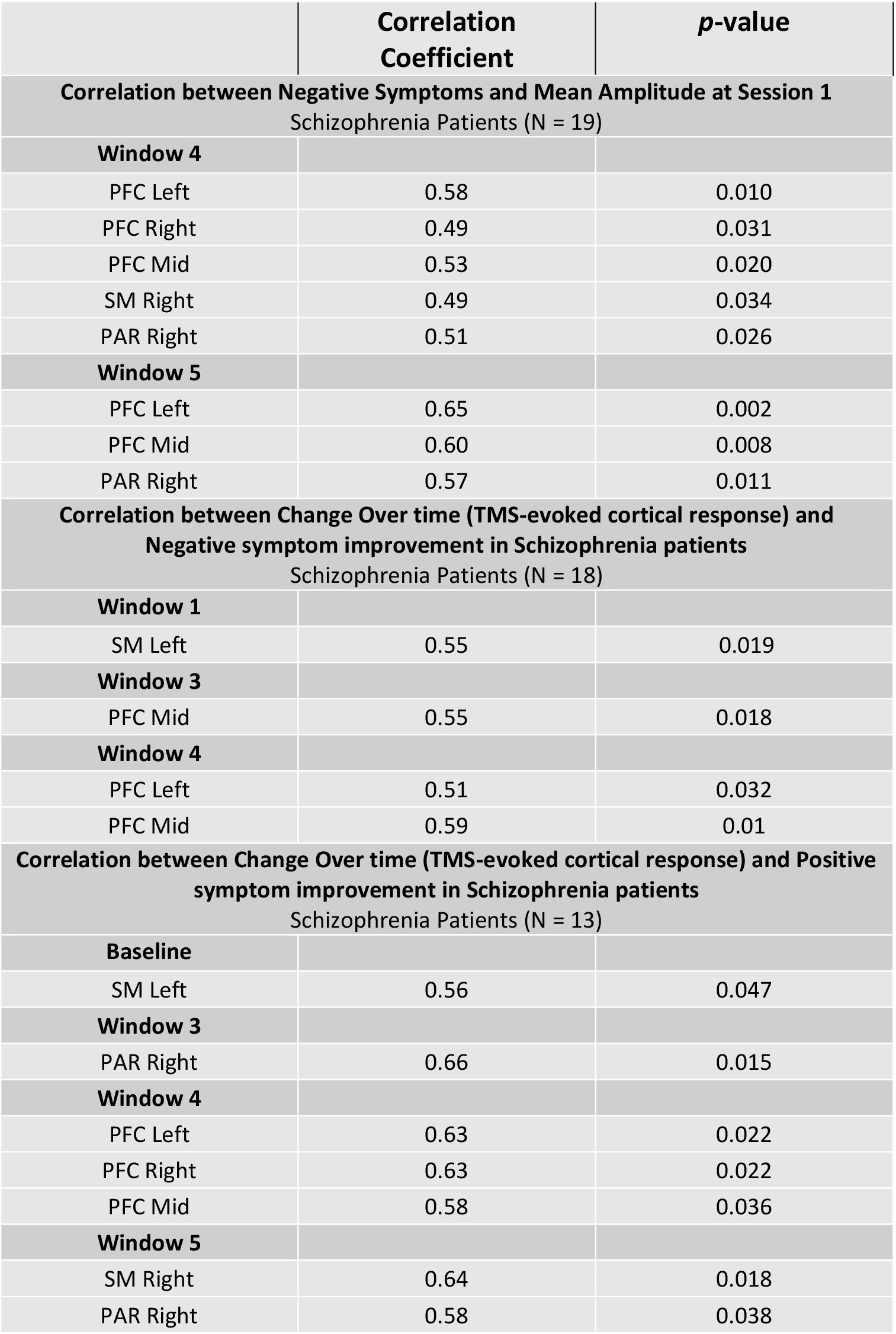
All significant correlations between the clinical symptoms of schizophrenia (negative and positive) and mean amplitude squared at Session 1. This is followed by all significant correlations between clinical symptoms of schizophrenia (negative and positive) and change in cortical response to TMS stimuli across sessions.

|  | **Correlation Coefficient** | ***p*-value** |
| --- | --- | --- |
| **Correlation between Negative Symptoms and Mean Amplitude at Session 1**<br>Schizophrenia Patients (N = 19) |  |  |
| **Window 4** |  |  |
| PFC Left | 0.58 | 0.010 |
| PFC Right | 0.49 | 0.031 |
| PFC Mid | 0.53 | 0.020 |
| SM Right | 0.49 | 0.034 |
| PAR Right | 0.51 | 0.026 |
| **Window 5** |  |  |
| PFC Left | 0.65 | 0.002 |
| PFC Mid | 0.60 | 0.008 |
| PAR Right | 0.57 | 0.011 |
| **Correlation between Change Over time (TMS-evoked cortical response) and Negative symptom improvement in Schizophrenia patients**<br>Schizophrenia Patients (N = 18) |  |  |
| **Window 1** |  |  |
| SM Left | 0.55 | 0.019 |
| **Window 3** |  |  |
| PFC Mid | 0.55 | 0.018 |
| **Window 4** |  |  |
| PFC Left | 0.51 | 0.032 |
| PFC Mid | 0.59 | 0.01 |
| **Correlation between Change Over time (TMS-evoked cortical response) and Positive symptom improvement in Schizophrenia patients**<br>Schizophrenia Patients (N = 13) |  |  |
| **Baseline** |  |  |
| SM Left | 0.56 | 0.047 |
| **Window 3** |  |  |
| PAR Right | 0.66 | 0.015 |
| **Window 4** |  |  |
| PFC Left | 0.63 | 0.022 |
| PFC Right | 0.63 | 0.022 |
| PFC Mid | 0.58 | 0.036 |
| **Window 5** |  |  |
| SM Right | 0.64 | 0.018 |
| PAR Right | 0.58 | 0.038 |

Figure 5A shows the locations at which increased change in cortical response was related to symptom improvement of negative symptoms in SCH patients. Negative symptom improvement positively correlated with increased change in the sensory-motor regions at the early window (Window 1) and with the frontal regions at the later windows (across Window 3 and 4). Figure 5B displays the locations at which increased change in cortical response was related to symptom improvement of positive symptoms in SCH patients. Positive symptom improvement correlated with increased change in cortical response across the sensory-motor regions region at baseline and increased change across the parietal, frontal and sensory-motor regions at the later windows (across Window 3, 4 and 5) (see Figure 5. and Table 3.).

**Figure 5.**
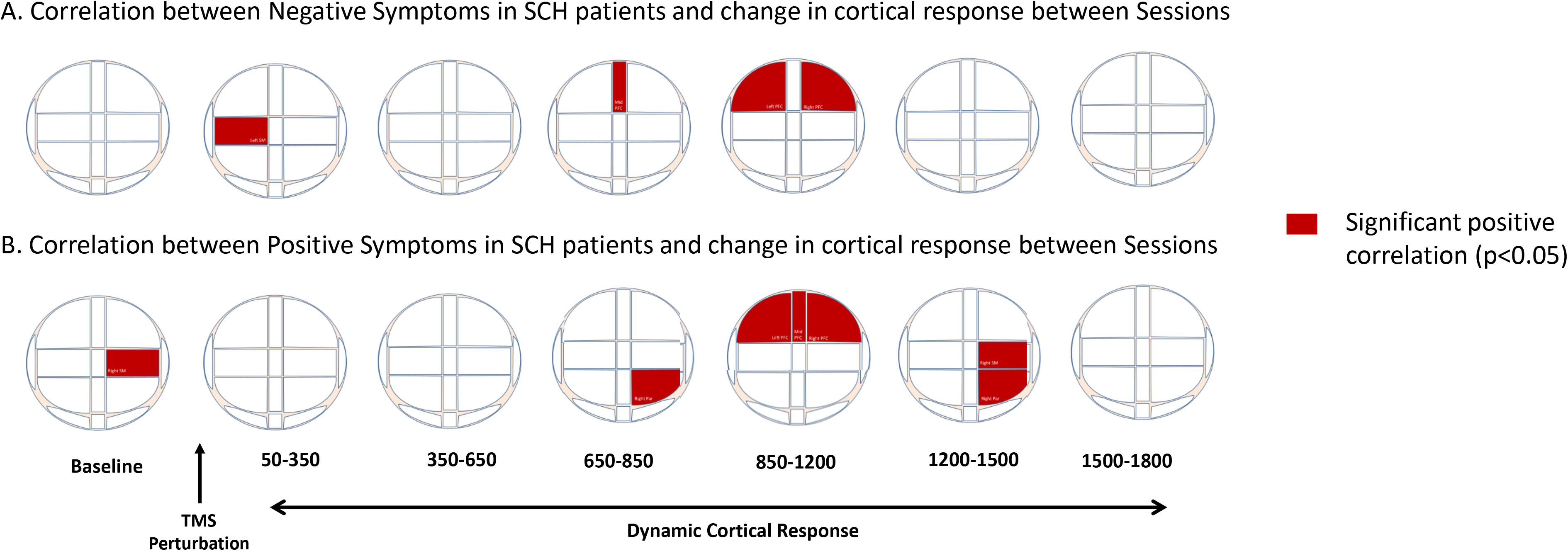
Locations of Significant correlations between change in TMS-evoked cortical response (between Session 1 to Session 3) and symptom improvement in schizophrenia (SCH) patients. **Figure 5A.** Demonstrates the locations of significant correlation between the increased change in TMS-evoked potentials over sessions and negative symptoms of SCH. **Figure 5B.** Illustrates the regions of significant correlation between increased change in TMS-evoked potentials over sessions and positive symptoms of SCH. Significance values are set at an alpha value of *(p* < 0.05) and only positive correlations were observed.

## Discussion

The current study identified abnormal patterns of cortical activity underlying stimuli processing in SCH patients. SCH presented with reduced excitability at the early time window followed by increased excitability at the later time windows compared to HC. The temporal sensitivity of this cortical response was only evident when the extended time-course of response was examined. Electrophysiological information extracted from later time windows was clinically-relevant and related to symptom severity in SCH patients. These patterns of cortical response vary more over sessions in SCH patients relative to HC.

Increased change in the cortical response over sessions in SCH patients was associated with improvement of SCH symptoms. This is the first study, to the authors knowledge, to have identified a clinical relationship between the reproducibility of the dynamic patterns of TMS-evoked cortical response in treatment-receiving SCH patients and symptom improvement.

### Dynamic patterns of cortical excitability underlying stimuli processing

In response to the frontal TMS stimuli, SCH patients showed *reduced* activation in the sensory-motor region in the early window and with *increased* activation in the frontal (local regions) and parietal (more remote) regions across the later time windows compared to HC. By extending the time-course of analysis, our study demonstrated the clinical importance of mapping the temporally-sensitive patterns of TMS-evoked cortical response in more adequately capturing the cortical processes underlying abnormal stimuli processing in SCH patients.

The majority of the earlier TMS-EEG studies in SCH patients focused on the early cortical response to TMS stimuli [14,15,20,22]. The early response to TMS may reflect a range of multisensory processes (including an integrated network of somatosensory, auditory and attentional resources activated by the stimuli) [37–39]. Consistent with the earlier findings, we observed reduced cortical response in the sensory-motor regions to the TMS stimuli at the early time window. The sensory-motor region is recruited by the frontal thalamocortical circuitry [40], a core network of the brain which mediates the bidirectional cortico-cortical information flow and cortical communication with the cerebral cortex [41] and is compromised in SCH patients [42–44]. Thus, the attenuated early activity within the sensorymotor regions may reflect disturbances in the integrity of connections between the brain regions involved in low-level sensory processing [45]. This concept is consistent with earlier TMS-EEG [14,20] and TMS-fMRI [46] studies which observed reduced thalamic activation in response to TMS stimuli in SCH patients. Therefore, we propose that dampened activity within the sensory-motor regions may reflect an inability to adequately process the early TMS-specific sensory information in SCH patients [47].

Following the early windows, SCH patients presented with widespread increased cortical activation across the frontal and parietal regions relative to HC before returning to comparable baseline levels. Increased excitability at these later windows is associated with more severe negative symptoms of SCH. This difference between SCH and HC is not surprising, as altered functioning across these higher-order regions have been widely implicated by neurobiological models of SCH [48] and underlie key cognitive features of SCH [49]. Moreover, the hyperexcitability of response at the later windows may be due to an impaired processing of early multisensory information (elicited by the TMS pulse) by SCH patients. Whereby, the ability to adequately inhibit and filter incoming irrelevant sensory stimuli is a fundamental protective mechanism that prevents flooding higher-order cortical areas with unnecessary information [7,8]. Disturbances in the processes underlying the filtering of inappropriate sensory stimuli has been found to result in neuronal hyperexcitability in SCH patients due to impaired neural inhibition at the subcortical and cortical levels [4,50]. Therefore, increased engagement of the higher cortical areas (such as the frontal cortex and the more remote interconnected parietal region) may be related to enduring deficits in the early processing (filtering and integration) of the TMS stimuli in SCH patients [23], with this effect being most prominent in patients with more severe symptoms of SCH.

### Test-retest: Variability in dynamic patterns of cortical excitability over sessions

Following the identification of the altered patterns of cortical response to TMS stimuli in SCH patients, we sought to determine whether these patterns remain stable over sessions or are amenable to change as symptoms of SCH improve. Using a test-retest approach, we found that SCH patients showed increased variability and less stability of response compared to HC over repeated sessions. This is the first report of the utility of the test-retest TMS-EEG protocol to identify increased variability in cortical response within a psychiatric population of SCH patients while the cortical response in healthy controls appears to be a useful comparative ‘control’ condition [24–26].

In the current study, all patients were undergoing pharmacological treatment for SCH and the greatest symptom improvement occurred between Session 1 and Session 3. Therefore, variability in cortical response between these sessions were examined. We identified a relationship between an increased amplitude of change in the cortical response to TMS and symptom improvement. This relationship was observed across a dynamic time range (baseline, early and later time windows) and over a range of brain regions (frontal, sensorymotor and parietal). Therefore, an increased flexibility to change within the frontal thalamocortical circuitry during treatment was indicative of symptom improvement. This is consistent with a recent test-retest fMRI study which identified a relationship between changes in functional connectivity within the thalamocortical circuitry and an improvement in cognitive features of SCH following treatment [51]. Thus, characterising the amenability to change in cortical response over the sessions by the SCH patients (while HC remain stable) may contribute to the identification of potential neuro-biomarkers of symptom-improvement. As such, the present study provides the first report that TMS-EEG may be useful not only in quantifying neural abnormalities underlying TMS stimuli processing in psychiatric populations but also in using this information to identify potential neural biomarkers of clinical improvement in SCH patients [13].

### Limitations

Several potential limitations of the current study need to be considered. Firstly, while EEG is a neuroimaging tool with excellent temporal resolution, it has poor spatial resolution. Therefore, interpretation of the exact circuitry underlying these dynamic brain processes (beyond the surface cortical sites) must be considered with caution. Secondly, the current study was the first to compare the finding of increased cortical excitability at later time windows with baseline measures [23]. Increased excitability across the frontal (left) and parietal regions in SCH patients were observed at later time windows which were not evident at baseline and appears to reflect a true cortical response to the TMS stimuli by SCH patients. Notably though, there was greater variability in response by SCH patients which may reflect a constant background noise [52,53] and this should be taken into account when considering the overall findings of the study and may relate to the increased noise in the baseline time window in the topological map of the response. Finally, it is important to consider that all SCH patients were undergoing anti-psychotic drug treatment and we are unable to separate the potential effects of specific antipsychotic medications from our results. We examined the effect of medication type (atypical vs typical anti-psychotic medication vs combination of treatments) on the cortical response and observed no significant changes in the current findings. Therefore, our findings should be considered as a representative of an in-patient and medicated SCH population.

## Conclusion

To conclude, our study quantified the dynamic patterns of cortical response underlying TMS stimuli processing in SCH patients. By mapping the response over an extended time-course, we found that SCH patients exhibited attenuated excitability at the early window and increased excitability at the later windows before returning to baseline. Increased excitability at later windows was related to increased symptoms of SCH. We propose that these altered patterns of cortical response may relate to impaired sensory gating and an inability to adequately filter incoming sensory stimuli, which in turn, may lead to hyperexcitability of higher order cortical areas due to a lack of inhibition of irrelevant information. This is in line with neurobiological studies of SCH, which show altered functioning of the thalamocortical circuitry, the same region which is required for adequate filtering of sensory information. It was then shown that these patterns of cortical response appear to change in treatment-receiving SCH patients while the response in HC remained relatively stable over sessions. This is an exciting area of research which requires further evaluation (as we provided a preliminary observation of the patterns of variability). However, a strong relationship was identified between the increased amplitude of change in cortical response over sessions in SCH patients and symptom improvement. Therefore, the present study maps the temporally-sensitive cortical abnormalities underlying stimuli processing in SCH patients and provides the first report of these dynamic patterns being amenable to change as symptoms of SCH improve. It is anticipated that this research will contribute to neurobiological models of SCH and hopefully lead to the identification of clinical neuromarkers to improve current diagnosis and treatment models.

## Acknowledgements

Sincere appreciation is expressed to Netali Mor, Gily Ginosar, Dana Koresh and Shir Yerushalmi for assisting with healthy control data acquisition and Dana Tel-Zur for her contribution in pre-processing the EEG data. Special thank you to the following IDC students. May Recanti, Yael Pugatch, Amit Ben-Zvi, Danielle Tabib, Hadar Glasberg, Liron Drori and Adi Harel for assisting with the patient data acquisition and Jackie Shpilman, Lia Hadas, Danit Rose, Aviv Sheriff, Zohar Goffen and Yael Spiegelman for their contribution in applying the TMS-EEG artifact correction and cleaning the EEG data. Dr. Jodie Naim-Feil is a recipient of the Senior Post-doctoral Fellowship at the Weizmann Institute, the Curwen-Lowy Post-doctoral Fellowship and the Clore Post-doctoral Fellowship which supported the development of this study. Prof. Elisha Moses is supported by the Minerva Foundation, Germany, and by the Israel Science Foundation. The study was supported by Dr. Nava Levit-Binnun’s Israel Science Foundation grant 1169/11 and by the National Institute of Psychobiology in Israel.

