## Supplemental Materials for "Variability of dynamic patterns of cortical excitability in schizophrenia: A test-retest TMS-EEG study"

### *Supplemental Information*

#### Electroencephalography (EEG) pre-processing steps

The continuous EEG data were segmented into epochs from -1000ms to 2000ms around the TMS stimuli. The EEG data was then baseline corrected (-1000ms to 0). TMS-induced discharge artifacts were removed using the procedure detailed by Freche et al (2018) (1). Data traces were filtered with a linear FIR bandpass filter (1-80Hz) and notch-filtered at 50Hz using EEGLAB's filter function (2). Epochs contaminated by extreme movement or muscle artifacts (such as yawning or facial twitches) were excluded via visual inspection by an EEG trained research assistant. To remove eye-blinks, lateral eye movement and auditory artifacts, we conducted Independent Component Analysis (ICA) and manually removed the three largest components containing these artifacts using EEGLAB (2).

### TMS-evoked potential (TEP) response to frontal (FC2) stimulation

A. TEP response to frontal TMS across 64 electrodes

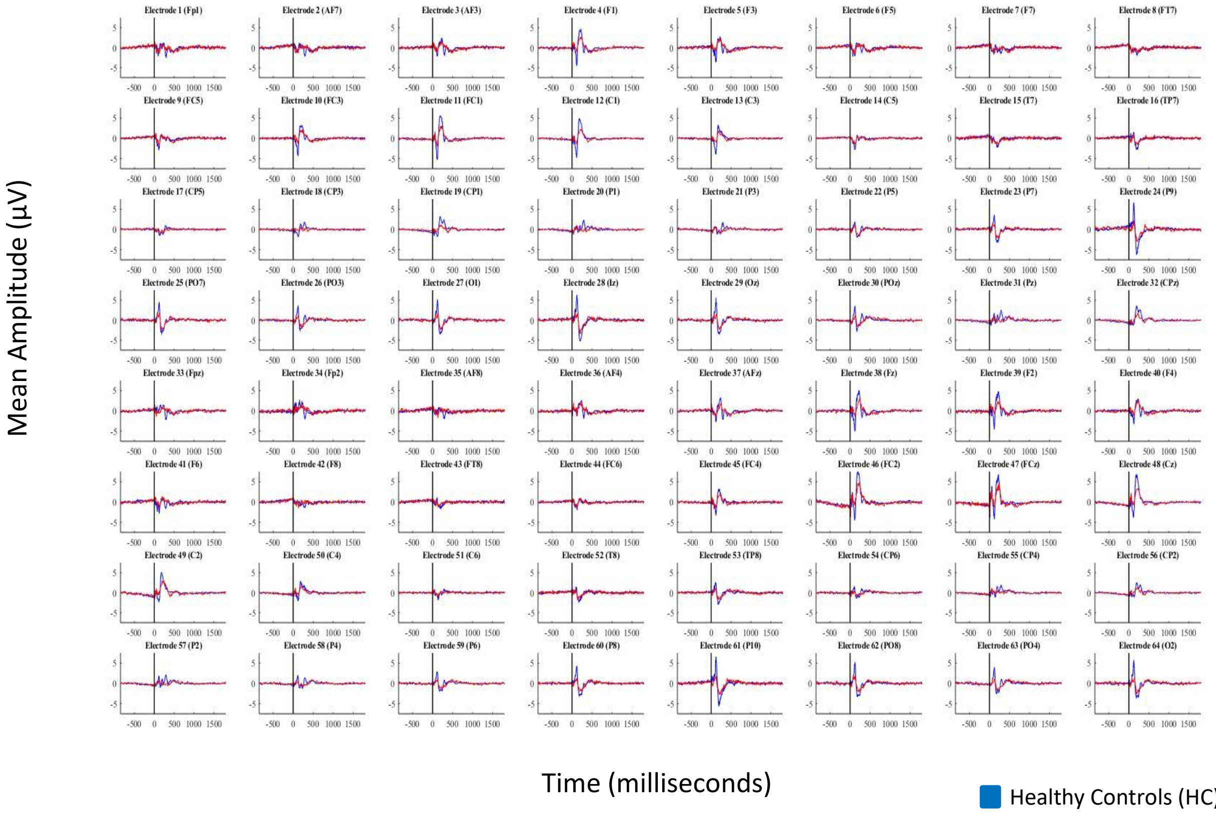


B. Magnified example of the TEP response to frontal TMS at electrode 4 (F1)

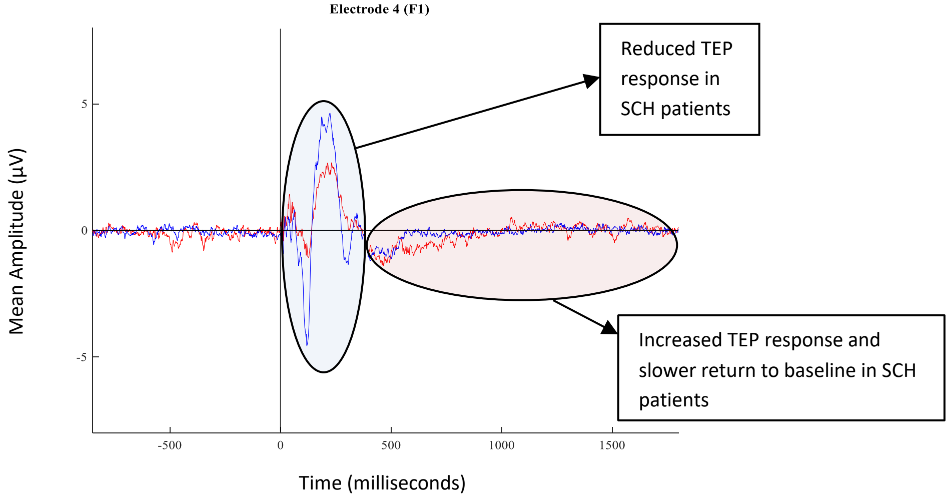


**Figure S1A.** illustrates the averaged TMS-evoked potential (TEP) response to 100 single transcranial magnetic stimulation (TMS) pulses applied to the frontal cortex (electrode FC2) across the 64 electrodes at Session 1 for schizophrenia (SCH) patients and healthy controls (HC). **Figure S1B.** magnifies the cortical response at electrode F1 and provides an example of the pattern of response observed across many electrodes. Whereby, SCH patients exhibit a dampened cortical response at the early time windows and increased cortical response and slower return to baseline at the later time windows.

**Division of 64 electrodes into 14 representative brain regions**

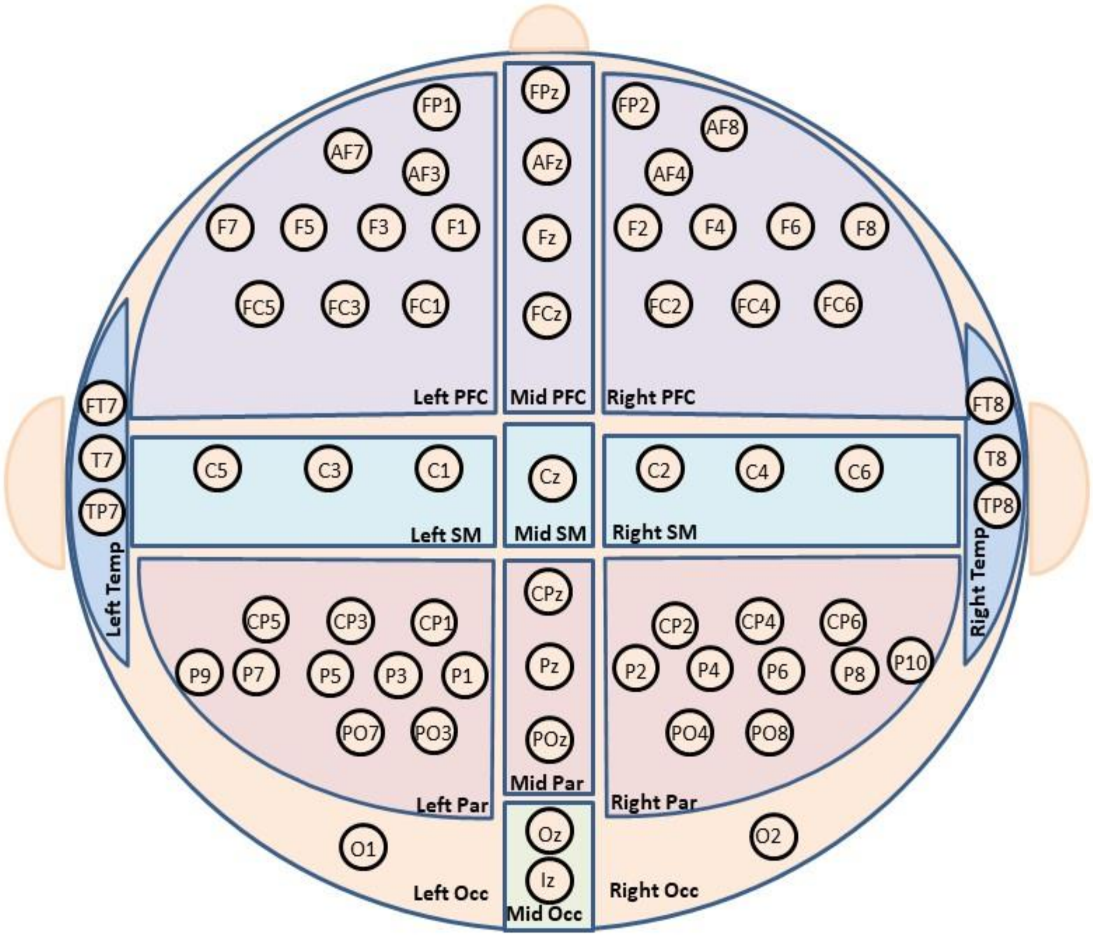


**Figure S2.** A schematic illustration of the division of 64 electrodes (according to the 10/20 International system) into fourteen representative brain regions (not drawn to scale). The TMS-evoked potentials were averaged over each of these brain regions for each participant. The division of these brain regions is based on the representative brain map detailed by Naim-Feil et al (2018) (3). Importantly, in the current study we do not make direct comparisons between regions (within groups), but rather, comparisons of the same regions across different groups (between groups). Therefore, the different number of electrodes averaged per region does not compromise the between group comparisons. Specific regions of interest were examined for this study (to examine the dynamic effect of the TMS-pulse) and these were defined as the PFC (left, right and Mid), SM (left and right) and the Par (left and right). PFC = prefrontal cortex; SM = Sensory-Motor; Par = Parietal; Occ = Occipital; Temp = Temporal.

### Standard Error of Measurement

The Standard Error of Measurement (SEM) provides a measure of 'spread' across the repeated measures (i.e. the three sessions) from the true score within stable individuals (4–6). The SEM calculates the standard deviation of all within-subject sources of variables (while excluding between-subject variances). In the current study, we aimed to extract whether there is increased variation in TMS-evoked potentials over the three sessions across the groups when no other significant changes have occurred. To do this, we extracted the Within Mean Square (using SPSS v.22) for each electrode as an estimate of the population variance. Figure S3. illustrates the SEM values for schizophrenia patients (SCH) and healthy controls (HC) across each electrode and for each time window (see Figure 5. of the main text for a topographical representation of these results). From observing the SEM values, it appears that SCH patients present with considerably increased SEM values compared to HC for many electrodes and across various time windows. This finding, while observational, provides initial evidence of an increased variability across the three sessions in SCH patients relative to HC. Following on from this observational finding, we next statistically quantified whether *change over time* (from Session 1 to Session 3) in SCH was significantly higher than in HC (described in the main text of the manuscript).

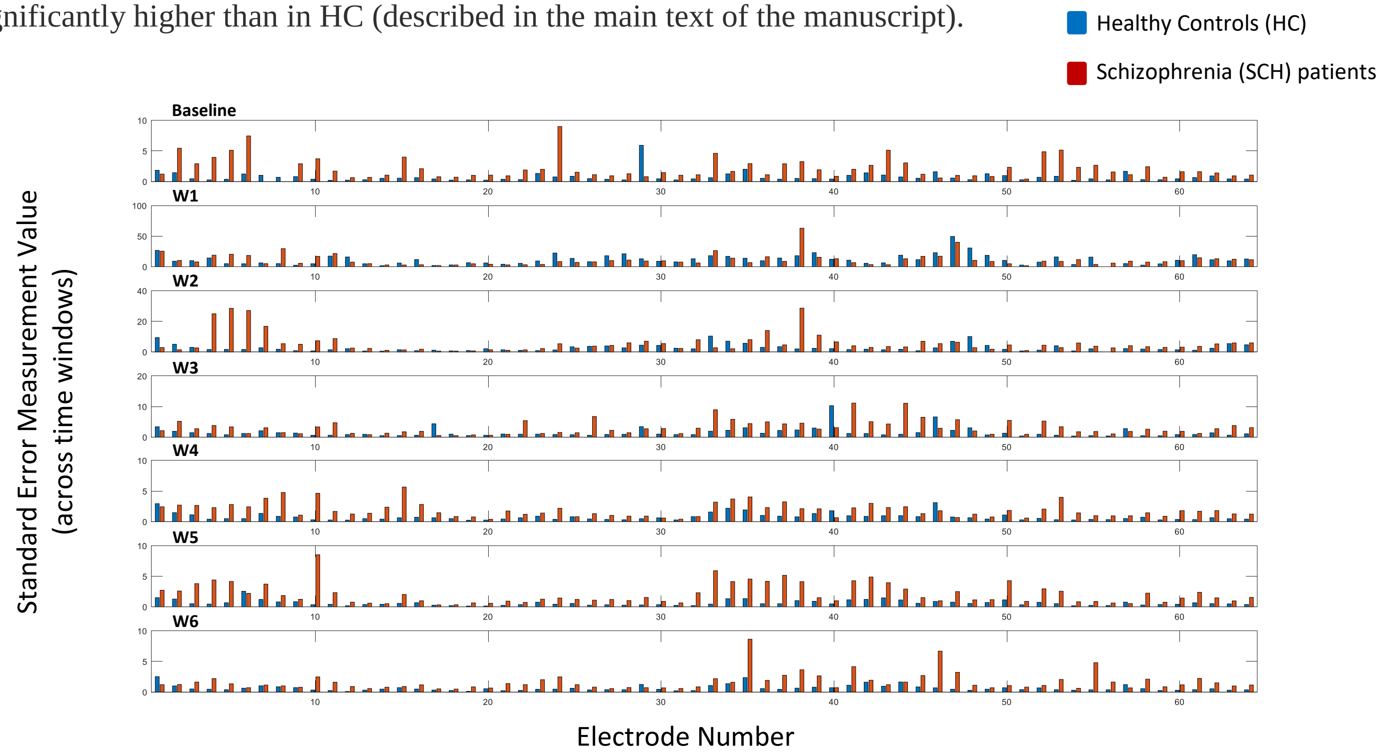


**Figure S3.** shows the Standard Error of Measurement (SEM) values across the 64 electrodes for schizophrenia (SCH) patients and healthy controls (HC). This provides initial evidence of increased variation in the TMS-evoked potential response of SCH patients over the three sessions compared to HC. The SEM values for all electrodes are presented (see Figure S1. for labels of electrodes) and across each time window (Baseline at -850ms to -550ms and W1 through W6 at 50-350ms, 350-650ms, 650-850ms, 850-1200ms, 1200-1500ms and 1500-1800ms respectively). Blue lines indicate healthy controls (HC) while red lines indicate schizophrenia patients (SCH).

### Cortical Response to TMS Stimuli averaged over the 7 representative brain regions

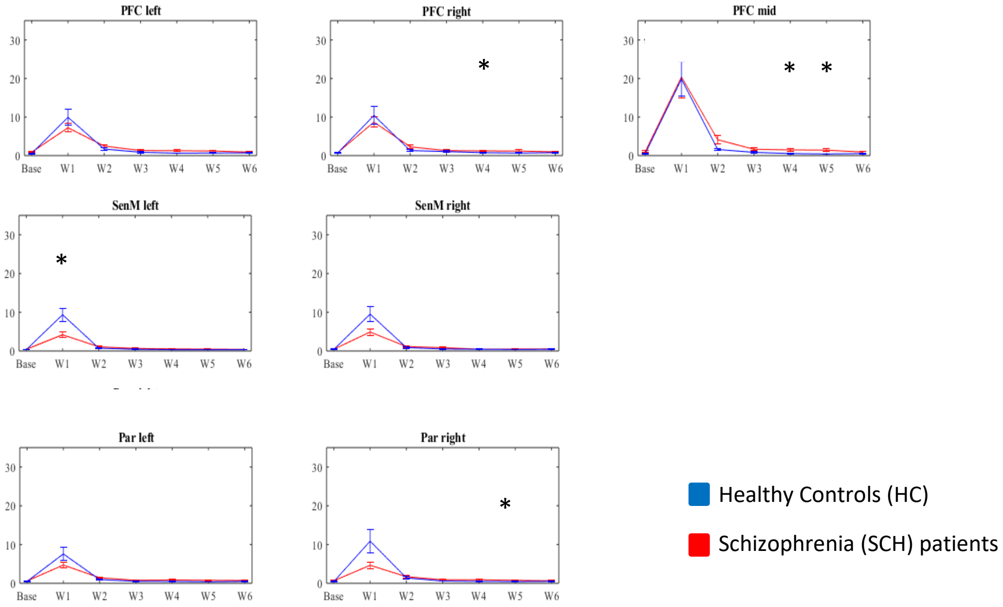


**Figure S4.** illustrates the TMS-evoked cortical response to the TMS stimuli averaged for each time window across the 7 brain regions for schizophrenia (SCH) patients and healthy controls (HC). SCH patients present with decreased activity in the early window (50-350ms) which is followed by increased activity in the later time windows (350ms onwards) in response to frontal TMS as well as a delayed return to baseline compared to HC. All significant differences between the groups are detailed in Table 1. of the main text of the publication. To illustrate the pattern of response we used a line graph format. It is important to note that the data is not continuous but rather averaged across each time window (Base at -850ms to -550ms and W1 through W6 at 50-350ms, 350-650ms, 650-850ms, 850-1200ms, 1200-1500ms and 1500-1800ms respectively). Blue lines indicate healthy controls (HC) while red lines indicate schizophrenia patients (SCH). * = $p$<0.05, ** = $p$<0.01. PFC = prefrontal cortex; SM = Sensory-Motor; Par = Parietal; Occ = Occipital; Temp = Temporal.
